# Altered Protein Phosphorylation in a Novel Midbrain Organoid Model for Bipolar Disorder

**DOI:** 10.1101/2025.04.01.646642

**Authors:** Katharina Meyer, Matthew Woodworth, Maria Carolina Bittencourt Gonçalves, Michelle Yue, Hoor AlJandal, Shad Morton, Michael Lewandowski, Ninning Liu, Eric Zigon, Patrick Fortuna, Mariana Garcia-Corral, Bogdan Budnik, George M. Church, Jenny M. Tam

## Abstract

Bipolar disorder (BD) is a severe psychiatric condition marked by episodes of mania and depression, with neurotransmitter imbalance in the midbrain believed to play a critical role in its pathophysiology. Despite this, there is currently no validated midbrain model for examining BD-associated molecular changes available. Leveraging recent advances in stem cell technology, we developed a midbrain organoid model using human induced pluripotent stem cells (hiPSCs) from BD patients and healthy controls (CTR). To address issues of variability and enhance the throughput in organoid production, we implemented liquid handling and high-content imaging techniques. Quality control metrics were established to identify organoids unsuitable for further study. Electrophysiological analysis via high-density microelectrode arrays (MEAs) revealed significantly elevated neuronal properties in individual BD organoids, including increased mean amplitude, conduction velocity, and extended axonal and dendritic growth. Transcriptome and proteome analyses indicated significant dysregulation of BD-relevant signaling pathways—such as those involving phosphatidylinositol, glycogen synthase kinase-3 beta, and AKT. Notably, we identified dysregulated casein kinase 2 (CSNK2A1) and calmodulin 3 (CALM3) in BD organoids, which were reversed by lithium treatment, highlighting potential novel targets for therapeutic intervention. This study validates the midbrain organoid model as a valuable tool for exploring the molecular underpinnings of BD and identifying new treatment avenues.

## Introduction

Bipolar disorder (BD) is a severe and debilitating psychiatric condition characterized by recurrent episodes of mania and depression^1^. Abnormal neuronal excitability and disrupted network dynamics have been consistently reported in BD, both in patient-derived neurons and animal models, suggesting altered electrophysiological properties as a core feature of the disorder^2,3^. For instance, studies have demonstrated impaired reward learning and altered feedback-related negativity (FRN) in manic and euthymic BD patients, indicating dysfunctional reward processing mechanisms^4^. However, many existing *in vitro* systems lack the cellular complexity or regional specificity required to accurately model complex network-level alterations^5^.

The development of region-specific brain organoids offers a promising platform to study such functional phenotypes in a more physiologically relevant context. Recent advances in stem cell technology have provided new avenues for investigating the underlying mechanisms of BD and paved the way for further utilizing brain organoid models to study cellular signaling events relevant to the disorder^6,7^. While the exact pathophysiology of the disorder remains elusive, one prominent hypothesis suggests persistent neurotransmitter imbalance within the midbrain contributing to the onset of both depression and mania^8–10^. The dopamine system in BD shows fluctuations linked to mood swings, with increased activity potentially contributing to mania and decreased activity potentially contributing to depression^8,9,11^ In fact, dysregulation of signaling pathways, including those involving brain-derived neurotrophic factor, extracellular signal-regulated kinases, and calcium signaling have been implicated in the development of mood disorders^12,13^.

To comprehensively explore the complex molecular landscape of BD, integrated omics approaches are increasingly being employed to map disease-associated changes across transcriptomic, proteomic, and phosphoproteomic levels^14^. This multi-layered analysis allows for the identification of coordinated alterations in gene and protein networks, uncovering signaling cascades and post-translational modifications that may contribute to pathophysiology. Moreover, these approaches enable the discovery of drug-responsive pathways and potential therapeutic targets by capturing the dynamic interplay between gene expression, protein abundance, and phosphorylation states in disease-relevant models^15^.

To date, no cellular midbrain model has been available to investigate the molecular changes associated with BD. To address this gap, in this study we developed and validated a region-specific midbrain organoid model derived from human induced pluripotent stem cell (hiPSCs) lines from 9 BD patients and from 11 healthy controls (CTR) with no neuropsychiatric disease history. By implementing automation and high-content imaging, we improved reproducibility and throughput of organoid generation. Using an integrated multi-omics approach (combining electrophysiological recordings, transcriptomics, proteomics, and phosphoproteomics), we characterized the functional and molecular landscape of BD midbrain organoids. Our findings reveal altered neuronal excitability, compensatory gene and protein expression changes, and lithium-responsive modulation of key signaling pathways, highlighting the value of this platform for uncovering disease mechanisms and therapeutic targets in bipolar disorder.

## Results

### Midbrain organoids derived from control and bipolar disorder hiPSC show similar structural properties and composition

Midbrain organoids derived from hiPSC mimic human midbrain structure and function, serving as crucial models for studying brain development, modeling diseases, and enabling drug discovery research^16,17^. Here we utilized existing protocols for generating midbrain organoids to gain insights into the neural mechanisms underlying BD^17^. Since organoid generation is a manual and labor-intensive process, it often is subject to variability. This variability can occur both between individual organoids and across different technical batches. One of the primary reasons for this inconsistency is the differences in handling techniques among various researchers. To reduce inconsistencies in cell composition, we introduced liquid handling techniques and high-content imaging. Specifically, we adapted and optimized the organoid workflow by incorporating the Fluent® 780 liquid handler (Tecan Trading AG, Switzerland) for automated media changes. In addition, we established midbrain organoid monitoring by high-content imaging using the multi-camera array microscope MCAM^TM^ (Ramona Optics, NC), which captures and analyzes wells with near-synchronous speed (Figure 1A).

**Figure 1:**
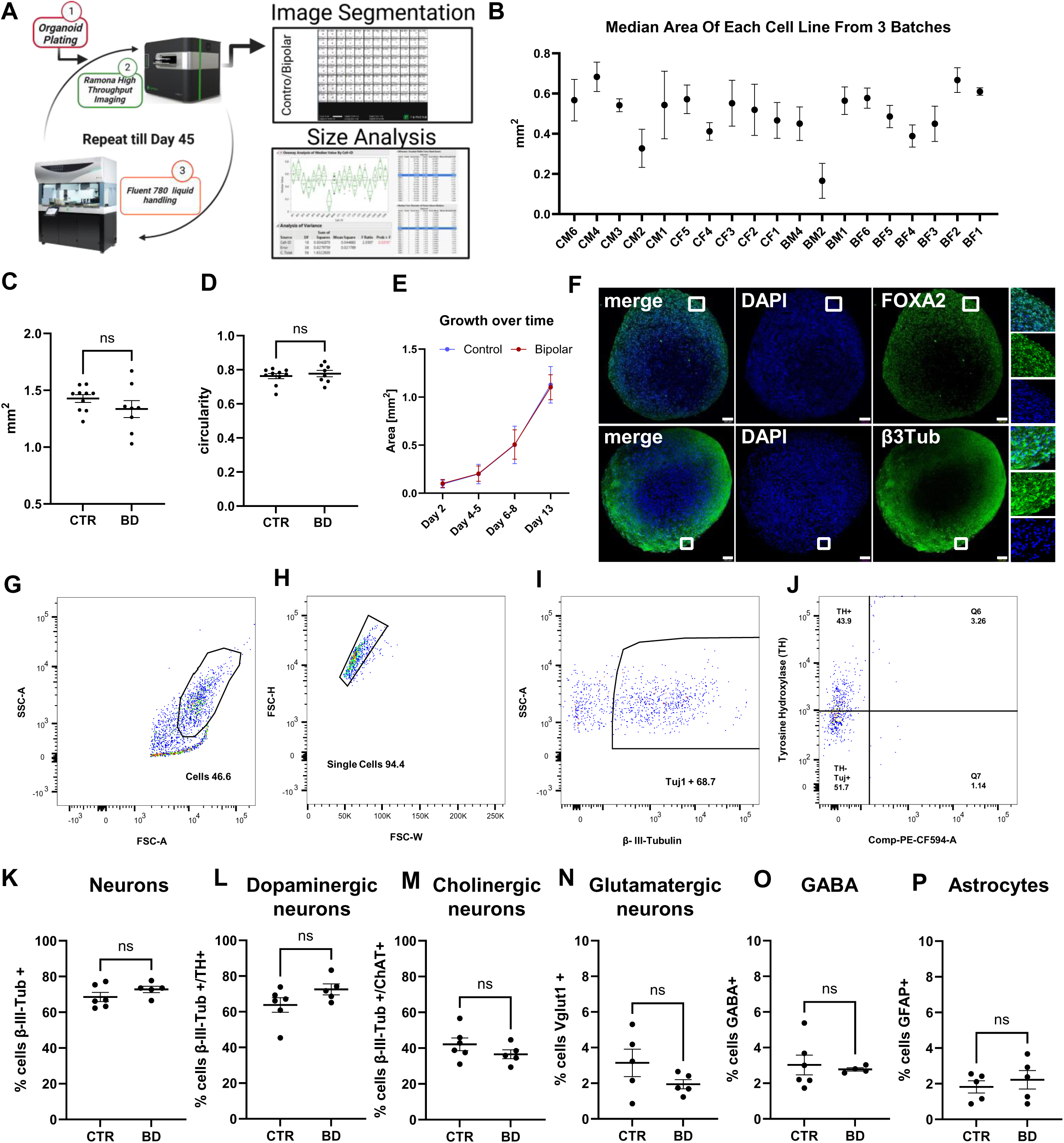
Comparable cell type distribution in CTR and BD Midbrain Organoids: **A** Schematic representation of semi-automated organoid quality control workflow. After embryoid bodies generation (1), organoids undergo repeated high-content imaging (2 – Ramona Optics) and liquid-handling assisted media changes (3 – Fluent 780). Resulting images are segmented and undergo size analysis. **B**: Median organoid area [mm^2^] calculated from 3 individual organoid batches at DIV7. **C-D**: Mean organoid area [mm^2^] and circularity in DIV35 midbrain organoids. Each datapoint represents the mean of 24 individual organoids from each hiPSC line. **E:** Mean area [mm^2^] of bipolar disorder (BD, n=8 with 24 organoids each) and **control** (CTR, n=8 with 24 organoids each) organoids plotted over time. **F**: Representative immunocytochemical staining of fixed midbrain organoids using FOXA2 (upper panel) and βIII-tubulin (lower panel) antibodies and DAPI. **G-J:** Representative gating strategies for FACS analysis of specific cell types. **K-P:** Quantification of cell type populations in BD (n=5) and CTR (n=6) using different marker antibodies: **K** Neurons (βIII-tubulin), dopaminergic (TH), cholinergic (ChAt), glutamatergic (Vglut1), GABAergic (GABA), astrocytes (GFAP). Data represent the mean ± SEM from CTR, n=5; BD, n=6. ***P<0.001 by Student’s t test.

To understand early developmental differences between organoids and establish early quality control metrics, we first generated three technical batches of embryoid bodies in 96-well plates. Subsequently, organoids from 8 control (CTR) and 8 bipolar disorder (BD) lines were repeatedly imaged until day *in vitro* (DIV) 35. Organoid area, circularity, and eccentricity were calculated to determine variation in growth for the identification of nonconform organoids early in the differentiation process. We observed that area and circularity are less variable than eccentricity between individual organoids, indicating that although organoids form circular structures, the deviation from a perfect circle towards more elliptical shapes is significant (Figure S1A-C). Consequently, eccentricity may not serve as a reliable metric for assessing the quality of organoid growth.

Following initial growth variations at early timepoints, we conducted an outlier analysis of organoid area and circularity at DIV7/8 (Figure S1D, E). By applying the ROUT outlier analysis with a Q value set at 5%, we could accurately identify organoids that significantly deviate in terms of circularity and area within their respective groups^18^ (Figure S1D, E). Additionally, we assessed the mean values for area and circularity across three independent organoid preparation batches and found that at least one line, BM2, failed to consistently produce satisfactory batches (Figure 1B and S1F). Although the differences from organoid batches of other lines were not statistically significant, this hiPSC line can be flagged for additional quality control assessments. Overall, we established a methodological approach to identify individual organoids that should be excluded from further analysis and to highlight specific hiPSC lines that may require further optimization and monitoring for consistent performance. Aiming to determine whether organoid growth is impaired in BD, we assessed growth metrics of CTR and BD organoids over time during early development (from DIV2-DIV13) and after completing the differentiation into midbrain organoids (at DIV35). Over a 10-day period, no significant differences in growth, nor in area, circularity, or eccentricity were observed at DIV35 (Figure 1C, D, E and S1G). Thus, BD and CTR organoids exhibited similar growth patterns and overall tissue structure.

To confirm neuronal identity and regionality of midbrain organoids, we performed immunostaining using antibodies against the midbrain floor plate marker, forkhead box A 2 (FOXA2) and βIII-tubulin (Figure 1F). Both markers are highly expressed in our model, confirming midbrain regional identity. To evaluate cell type distribution between BD and CTR organoids, we performed FACS sorting using antibodies against pan-neuronal (βIII-tubulin), dopaminergic (TH – tyrosine hydroxylase), cholinergic (ChAt – choline acetyltransferase), GABAergic (GABA), and glutamatergic (VGlut1 – vesicular glutamate transporter 1) neuronal markers (Figure 1G-J and S1 H-L). In addition, we assessed the presence of astrocytes using the marker GFAP. We found an overall population of neurons of about 65-75% in both CTR and BD (Figure 1K). Further subtype stratification revealed that dopaminergic neurons comprise approximately 60-70% of the population, while cholinergic neurons account for 35-40% (Figure 1L, M and S1M). Additionally, we observed a minimal presence of glutamatergic and GABAergic neurons, each constituting around 1-2% of the population (Figure 1N, O). Astrocytes consistently represented about 2% of the cells in both CTR and BD contexts (Figure 1P). This data indicates that control and bipolar organoids exhibit a remarkably similar distribution of cell types.

### Midbrain organoids from BD patients exhibit increased neuronal function

Neuronal hyperactivity has been observed in BD cortical organoids and altered neural network activity in neurons derived from BD hiPSCs. This suggests that neural network instability and hyperexcitation linked to mood instability in BD can be effectively modeled using hiPSC-derived neurons and cerebral organoids^7,12^. We evaluated the electrophysiological properties of individual neurons and neuronal networks using high-density MEA recordings and spike sorting algorithms to determine whether a similar phenotype exists in midbrain organoids.

Initially, three organoids, matured to DIV 51, were positioned and adhered to the MEA plate (Figure 2A). After a short acclimation period, we first performed an activity scan of both CTR and BD organoids (Figure 2B-G). The scans revealed no significant differences in firing rate (Figure 2B, C, H) amplitude (Figure 2D, E, I), or inter spike interval (Figure 2F, G, J) between BD and CTR organoids, indicating no detectable differences in neuronal network activity. We then measured electrophysiological activity from individual neurons using axon tracking software (Figure 2K, L). Strikingly, neurons in BD organoids showed a significant increase in mean amplitude at the initiation site, suggesting stronger depolarization and more efficient action potential firing (Figure 2M). In addition, the mean conduction velocity was significantly higher in BD neurons, suggesting altered signal transmission (Figure 2N). Importantly, the structural quality of neurons was altered in BD, with mean total axon length (Figure 2O), longest branch length (Figure 2P) and mean longest distance to initiation site (Figure 2Q) being significantly greater, suggesting structural remodeling and growth, potentially in response to more synaptic input. Overall, these results indicate a hyperexcitability phenotype in midbrain neurons of BD patients-derived organoids, consistent with previous observations in other hiPSC-derived neural models of the disease.

**Figure 2:**
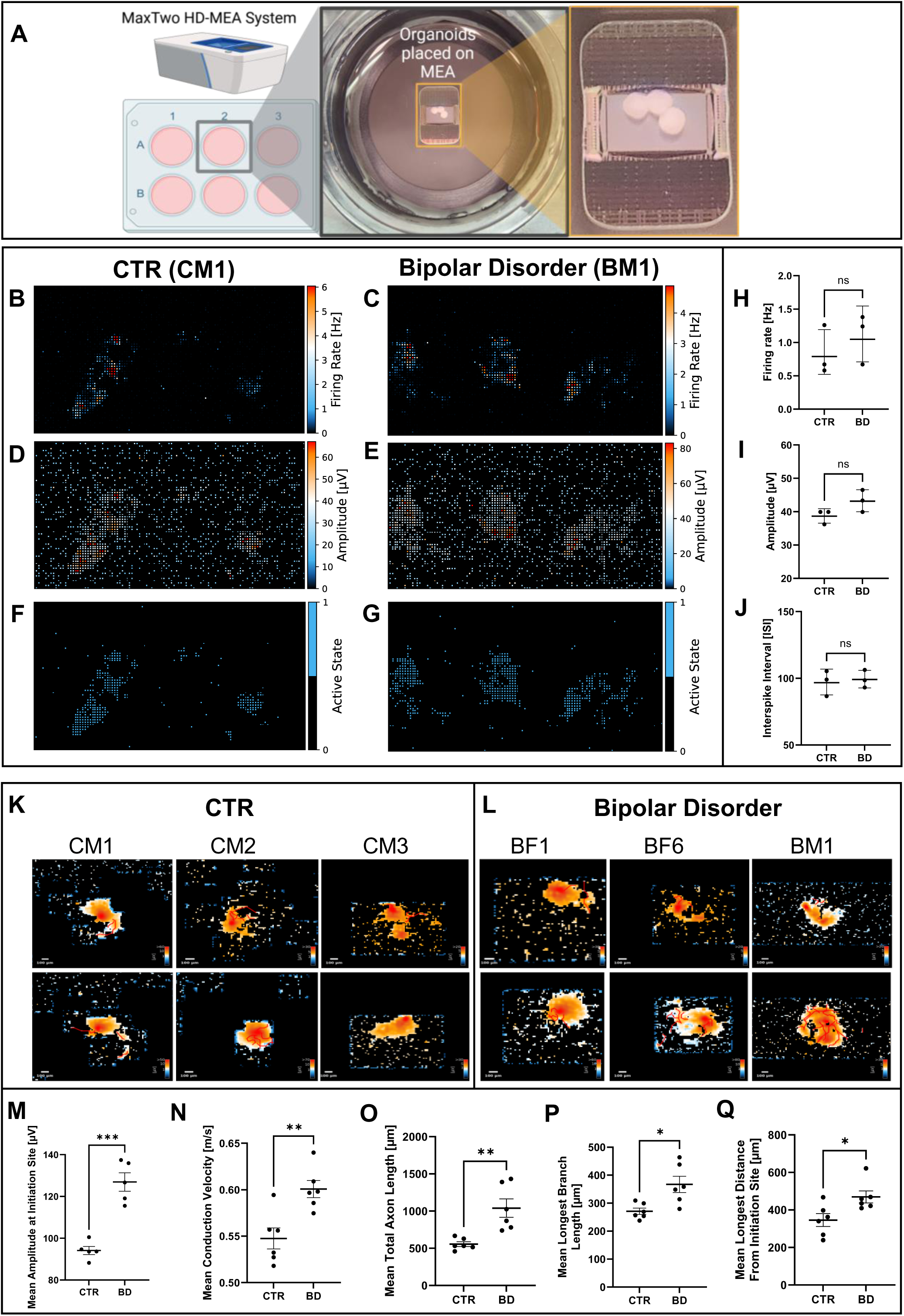
Increased neuronal function in BD midbrain organoids: **A** Schematic representation of microelectrode array (MEA) system and organoid placement on the electrode array. **B-G**: Activity assay-derived electrical activity. Representative images of activity maps showing firing rate (B-C), amplitude (D-E), active state (F-G) in one representative control (CTR, CM1) and bipolar disorder (BD, BM1) organoid line. **H-J**: Quantification of firing rate (H), Amplitude (I) and Interspike Interval (J) in BD vs CTR organoids. **K-L:** Representative images of axon tracking of individual neurons in 3 BD and 3 CTR lines. **M-Q**: Quantification of neuronal activity from individual neurons showing Mean amplitude at initiation site (M), mean conduction velocity (N), mean total axon length (O), mean longest branch (P) and mean longest distance from initiation site (Q). Data represents the mean ± SEM from several neurons from n=3 CTR and n=3 BD organoid lines (3 organoids each) measured over 6 individual time points (DIV49-56) ***P<0.001 by Student’s t test.

### Transcriptome changes suggest compensatory gene regulation in response to hyperexcitation

To uncover underlying mechanisms leading to the hyperexcitability phenotype, we first performed bulk transcriptome analysis from 5 CTR and 4 BD lines. The PCA plot shows a clear group separation of BD and CTR organoids (Figure 3A, Table S1). Hierarchical clustering reveals clear transcriptional differences between CTR and BD (Figure 3B). We then performed gene enrichment analysis using Gene Ontology databases (Figure 3C). Unexpectedly, we found down regulated genes to be associated with anion and cation channel activity pathways, while upregulated pathways included calcium ion binding and kinase activity – which is in stark contrast with the excitability phenotype and structural differences measured by MEA. These findings suggest the presence of a compensatory gene regulation network in BD organoids.

**Figure 3:**
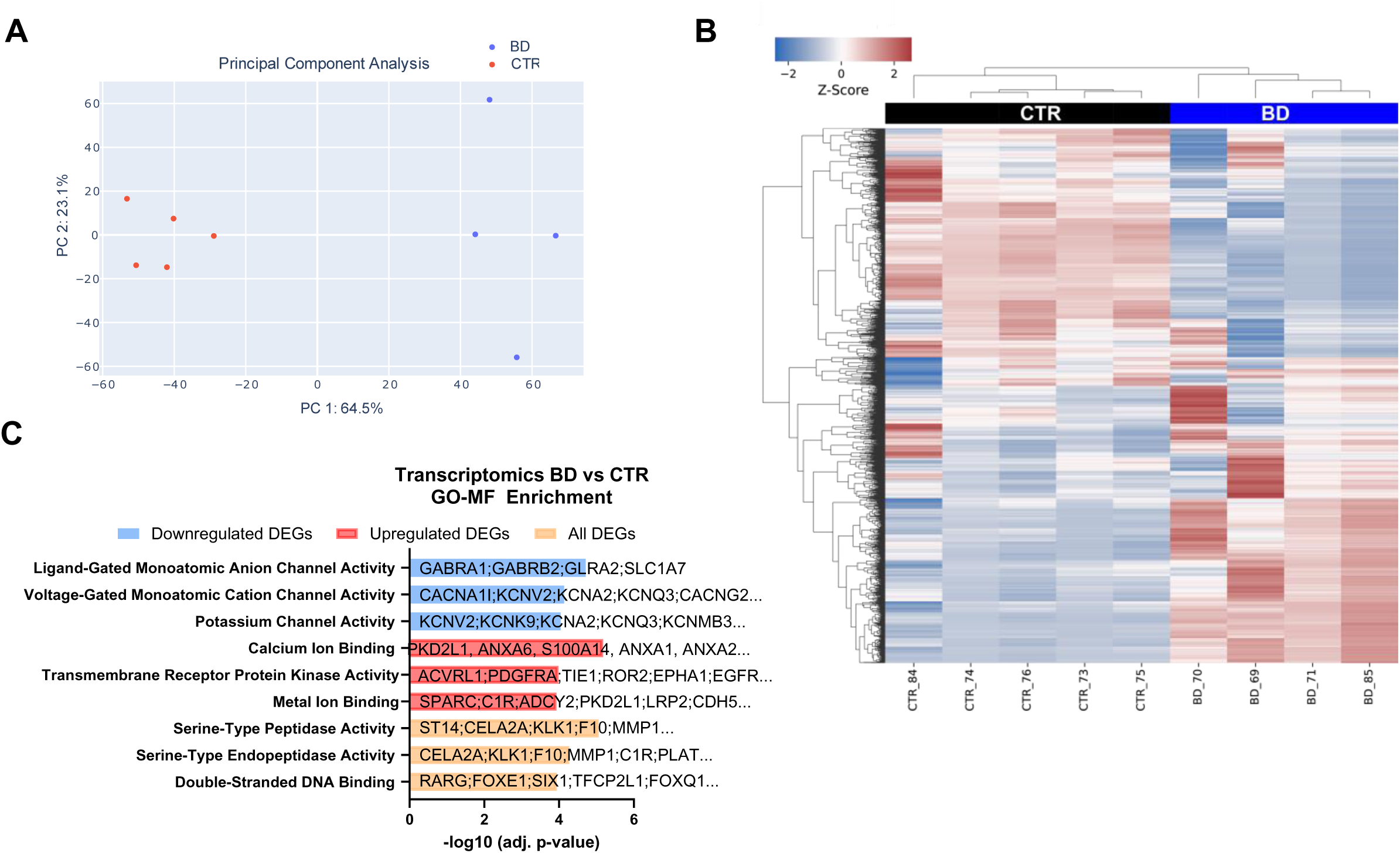
Transcriptome analysis of BD organoids reveals compensatory gene network: **A** Principal component analysis of bulk transcriptome data showing control (CTR, n=5) and bipolar disorder (BD, n=4) organoid sample distribution and clustering. **B**: Unsupervised hierarchical clustering of genes differentially expressed between CTR and BD. **C:** GO molecular function (GoMF) groups enriched in differentially expressed genes in BD compared to CTR organoids.

To further investigate protein-level changes, we performed unbiased proteomic analysis of midbrain organoids derived from BD hiPSC lines. Organoids from the same lines that underwent RNA sequencing were collected for TMT labelled mass spectrometry. We found a total of 234 significantly differentially expressed proteins (P<0.05) (Figure 4A, Table S2). Gene ontology analysis of pathways encompassing molecular functions, biological processes, and cellular components revealed that proteins associated with carboxylic acid binding, lysophospholipase activity, and spine morphogenesis within dendritic and axonal compartments are among the most upregulated hits (Figure 4B-D). These processes are consistent with the increased neuronal activity observed in our BD midbrain organoids. Notably, proteins that are downregulated predominantly fall into categories related to signaling, including adenylate cyclase activity, NADPH binding, and protein autophosphorylation (Figure 4B-D). These processes are crucial for the integration of neuronal receptor activation with intracellular signaling pathways. Specifically, we observed a significant downregulation of calmodulin isoform 3 (CALM3) and an increase in casein kinase 1 (CSNK1A1) expression (Figure 4E-F). Calmodulin acts as a critical mediator in calcium signaling and is essential for the regulation of CaMKII activity^19,20^, while casein kinases regulate circadian rhythms through phosphorylation of clock proteins and play central roles in Wnt/β-Catenin signaling cascades^21,22^. Disruptions in these pathways are implicated in the pathophysiology of bipolar disorder, affecting mood regulation and clinical symptoms^23–28^, raising the question of whether dysregulation at the level of substrate phosphorylation is also present in BD midbrain organoids.

**Figure 4:**
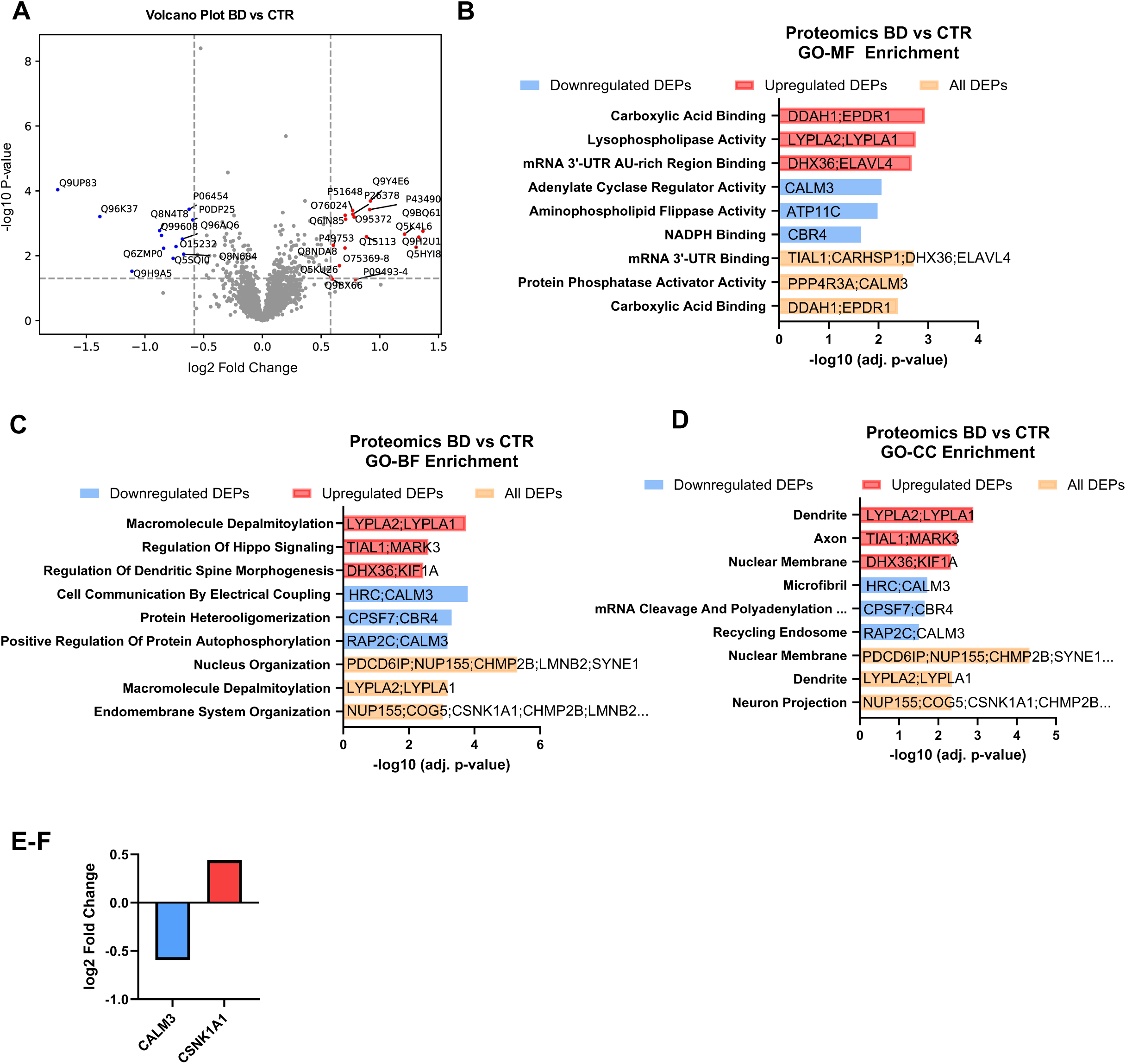
Bulk proteome analysis of CTR and BD Organoids: **A** Volcano plot showing differentially expressed proteins between bipolar disorder (BD) and control (CTR) midbrain organoids **B-D**: Gene Ontology molecular function (GoMF), biological pathways (GoBP) and cellular compartment (GoCC) groups enriched in differentially expressed proteins in BD compared to CTR organoids. **E-F:** Protein abundance of CALM3 and CSNK1A1 in BD vs CTR.

Phosphoproteome is largely impacted in BD and shows dysregulation of calmodulin and casein kinase substrate activation.

We performed unbiased phosphoproteomics analysis of midbrain organoids derived from BD (n=3) and CTR (n=3) hiPSC lines. We found a total of 196 differentially phosphorylated proteins in BD when compared to CTR (Figure 5A, Table S3). As a result of gene ontology enrichment analysis hyper-phosphorylated proteins clustered in pathways associated with the nuclear compartment and the regulation of RNA processing, while hypo-phosphorylated proteins clustered in pathways associated with the regulation of nervous system processes and the endoplasmic reticulum tubular network (Figure 5B). A string protein-protein interaction clustering analysis was performed on both hypophosphorylated and hyperphosphorylated protein sets (Figure 5C-I) since gene ontology enrichment analysis might not be suitable to analyze phosphorylated protein sets.

**Figure 5:**
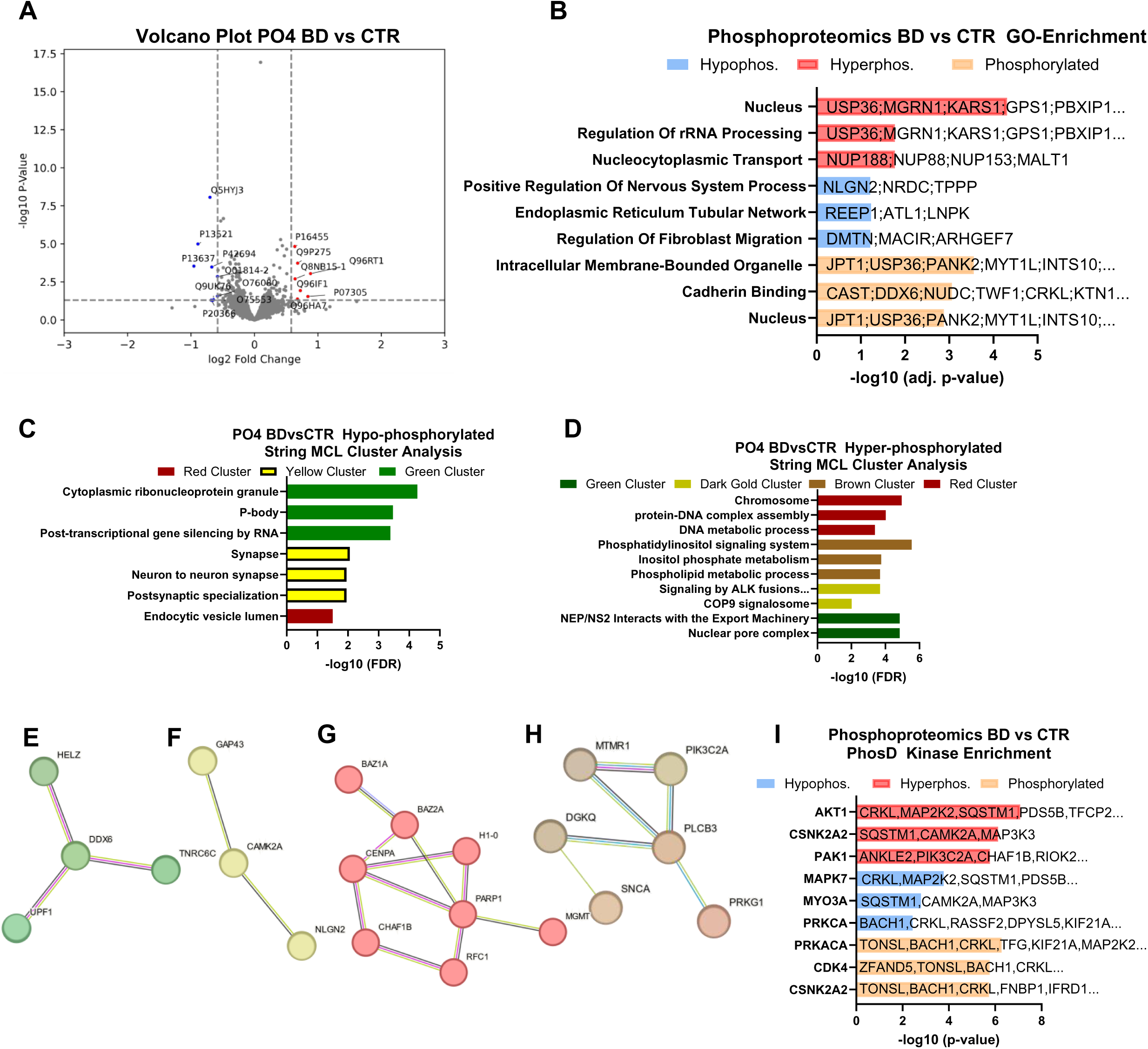
Bulk phosphoproteome (PO4) analysis of CTR and BD organoids: **A** Volcano plot showing differentially phosphorylated proteins in bipolar disorder (BD, n=3) and control (CTR, n=3) midbrain organoid lines. **B-D**: Gene ontology enrichment of hyper- and hypo- phosphorylated proteins. **C-D:** String protein-protein interaction network clustering of hypo- (C) and hyper-(D) phosphorylated proteins. **E-H:** Representation of clusters and proteins identified and shown in C-D. The hypo-phosphorylation clusters green and yellow (E, F) and hyper-phosphorylation clusters red and brown (G, H). **I:** PhosD kinase enrichment analysis of phosphorylated proteins.

A significant cluster of hypo-phosphorylated proteins was found in processes related to ribonucleoprotein granules, P-bodies, and RNA-silencing, indicating reduced post-translational activation and regulation. Hypo-phosphorylation was also observed in synapse and postsynaptic specialization processes, suggesting decreased activation of signal cascades. Notably, CAMK2A showed substantial hypo-phosphorylation, consistent with downregulation of CALM3 protein expression. In response to calcium activation calmodulins, including CALM3, bind to CAMK2A and induce a conformational change that allows autophosphorylation to occur. Furthermore, the observed reduction in phosphorylation of GAP43 and NLGN2 may alter synaptic function and neural plasticity by changing protein complex formation at synapses.

Hyperphosphorylated proteins, on the other hand, clustered in pathways that are important for protein-DNA complex assembly and DNA metabolic processes. Hyperphosphorylation of the proteins BAZ1A, BAZ2A, CENPA, H1-0, PARP1, CHAF1B, and RFC1 might upregulate processes such as chromatin remodeling and DNA repair to accommodate increased demands for gene expression and structural adaptations associated with synaptic plasticity (Figure 5D and 5G). Moreover, our data show that phosphatidylinositol (PI) signaling system is highly implicated in hyper-phosphorylated protein sets. Notably, disruptions in the PI signaling pathway – including altered inositol levels and protein kinase C (PKC) activity – have been linked to the pathophysiology of BD, and are direct targets of the mood-stabilizing drug lithium^29–31^ (Figure 5D, H).

To gain a deeper insight into the kinase-substrate relationship, we conducted kinase enrichment analysis with Kinase Enrichment Analysis 3 (KEA3), which incorporates both experimentally determined and predicted kinase-substrate interactions (KSI), as well as kinase-protein interactions (KPI), and interactions substantiated by co-expression and co-occurrence data^32^. When extracting the enrichment from the probabilistic kinase prediction model phosD^33^, we observed significant enrichment for AKT1, casein kinases and PAK1 (Figure 5I). Interestingly, AKT1, a key protein in the signaling pathway downstream of the PI system, has been implicated in the pathophysiology of BD, with genetic variations in AKT1 associated with increased risk for the disorder. Notably, this signaling pathway is also targeted by lithium^34–36^. Importantly, we identified enrichment of casein kinase CSNK2A1, and given the significant overlap between CSNK2 and CSNK1 targets, this aligns with the increased abundance of CSNK1A1 protein in BD organoids. Finally, we observed that the hypo-phosphorylation of CALM3 targets, consistent with reduced CALM3 protein levels, contrasts with the hyper-phosphorylation of CSNK1A1 targets, reflecting the increase in CSNK1A1 protein levels in BD midbrain organoids.

### Lithium treatment normalizes protein expression of calcium binding proteins and CSNK2A1/A2 kinase

The mood-stabilizer lithium is the first-line treatment for managing BD, and its neuroprotective and neurotropic effects are well documented in a variety of cellular models^37^. Several enzymes have been proposed as potential targets of lithium action, including inositol monophosphatase (IMP), a family of structurally related phosphomonoesterases, and the GSK3B^38^. Most of these targets are widely expressed, rely on metal ions for catalysis, and are inhibited by lithium via an uncompetitive mechanism, most likely by displacement of a divalent cation, such as magnesium^39^. Consequently, the landscape of lithium’s targets and mechanisms of actions remain complex and incompletely understood.

The midbrain organoid model offers a unique tool for accessing dynamic changes mediated by lithium treatment within a complex neural tissue environment. Thus, we explored whether lithium ameliorates effects on CALM3 and CSNK2A1/2 expression as well as kinase activation. First, we analyzed unbiased phosphoproteomes from CTR (n=3) and BD (n=3) hiPSC lines after treatment with 100 µM lithium carbonate (Li_2_CO_3_) for 1 week starting at DIV46. After lithium treatment, we observed differential phosphorylation profile in a total of 752 and 297 proteins in CTR and BD, respectively (Figure 6A, B, Table S4). As expected, lithium treatment had similar effects on the phosphoproteome in both BD and CTR organoids; however, only ∼75% of phosphoproteins were shared between the two groups, suggesting that 77 phosphoproteins are targeted by lithium uniquely in the BD context (Figure 6C). While CTR organoids showed a ∼25% overlap with BD, they retained 532 uniquely phosphorylated proteins following lithium treatment. The results of our study provide additional evidence for the importance of considering human disease backgrounds when studying drug effects^40^.

**Figure 6:**
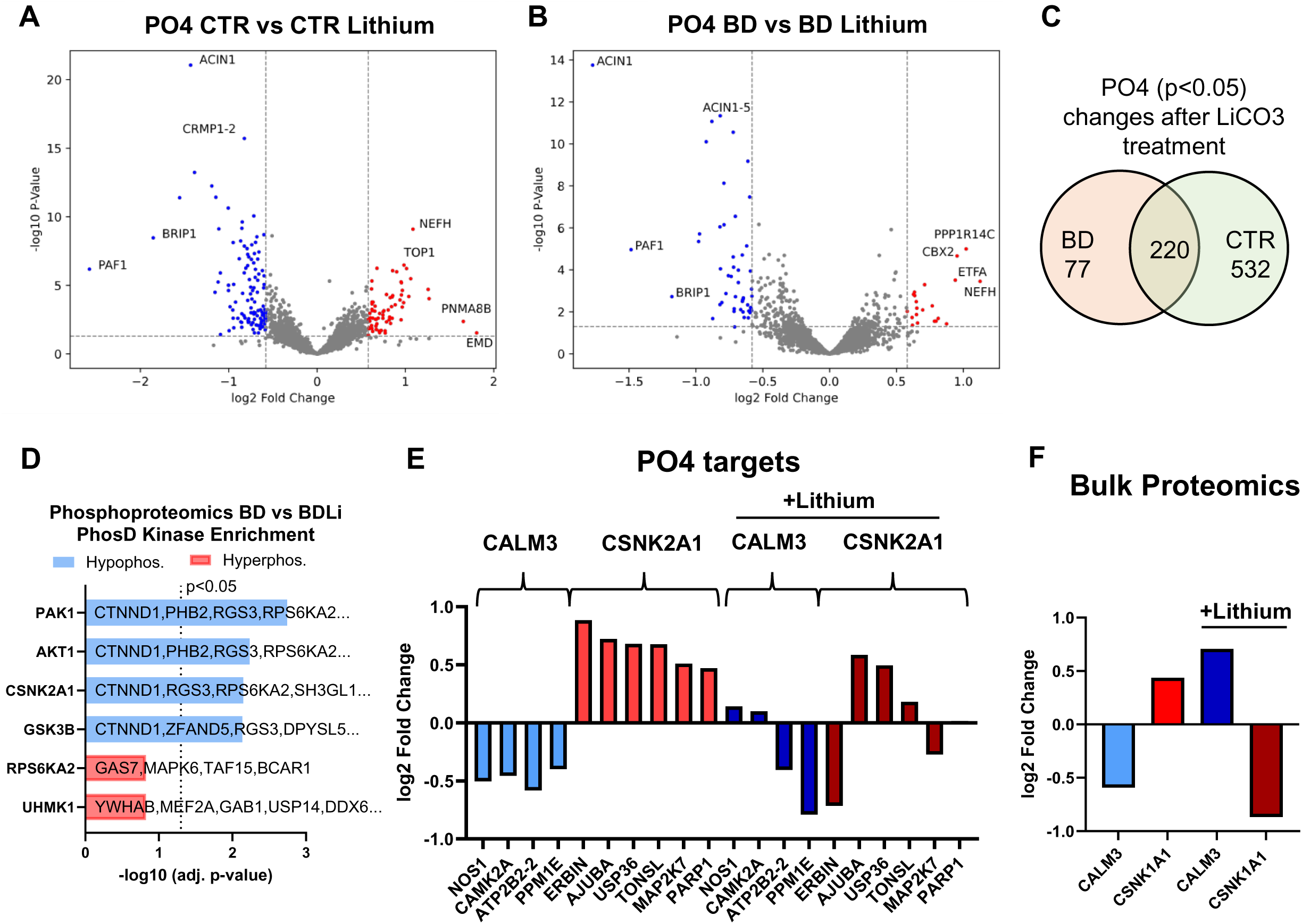
Bulk phosphoproteome (PO4) analysis of CTR and BD organoids following lithium treatment: **A-B** Volcano plot showing differentially phosphorylated proteins in bipolar disorder (BD, n=3 and control (CTR, n=3) midbrain organoid lines after treatment with 100µM Li_2_CO_3_ for one week. **C:** Venn diagram showing phosphorylated proteins overlap between CTR and BD after lithium treatment. **D:** PhosD kinase enrichment analysis of phosphorylated proteins in BD after lithium treatment. **E:** Plotting of log2FC of CALM3 and CSNK2A1 substrates before (BD vs CTR) and after lithium (BD vs BDLi) treatment. **F**: Protein abundance of CALM3 and CSNK1A1 before (BD vs CTR) and after lithium (BD vs BDLi) treatment.

We then performed kinase enrichment analysis to assess the effect of lithium on upstream kinases in BD midbrain organoids (Figure 6D). In line with previous findings, PAK1, GSK3B and AKT1 were significantly enriched among the hypo-phosphorylated protein sets. These findings further support the role of AKT1 and GSK3B as relevant nodes in lithium-responsive signaling networks^34–36,38,41^. In addition, PAK1 and AKT1 exhibit a reciprocal relationship, as PAK1 can either activate or be activated by AKT1^42,43^. These observations present an important point of validation of our midbrain organoid model. Importantly, we also found CSNK2A1 enriched among the hypo-phosphorylated protein sets. This contrasts with our earlier observations in BD versus CTR organoids, where CSNK2A1 was enriched in the hyper-phosphorylated set, suggesting a lithium-dependent reversal of CSNK2A1 activity (Figure 6D).

Kinase enrichment among hyper-phosphorylated proteins did not reach statistical significance. To examine this further, we compared the phosphorylation state of CALM3 and CSNK2A1 substrates between BD vs CTR as well as BD vs lithium-treated BD organoids, and observed a partial reversal (Figure 6E). Specifically, the hypo-phosphorilation of neuronal nitric oxide synthase NOS1 and CAMK2A was restored upon lithium treatment, while the plasma membrane calcium-transporting ATPase 2 (ATP2B2) and the protein phosphatase methylesterase (PPME1) remained hypo-phosphorylated. Similarly, CSNK2A1 substrates such as Erbb2 Interacting Protein (ERBIN), Tonsoku-Like, DNA Repair Protein (TONSL), Mitogen-Activated Protein Kinase Kinase 7 (MAP2K7), and Poly (ADP-Ribose) Polymerase 1 (PARP1) were no longer significantly hyper-phosphorylated following lithium treatment, whereas AJUBA and Ubiquitin-Specific Peptidase 63 (USP63) remained hyper-phosphorylated. (Figure 6E). A comparison of bulk proteomics data between BD and CTR and BD and BD after lithium exposure revealed a complete reversal in protein expression: CSNK1A1 was downregulated, while CALM3 was upregulated (Figure 6E, F, Table S5). Together, these findings demonstrate the utility of midbrain organoids for studying BD pathophysiology and lithium’s dynamic effects on the proteome and phosphoproteome. In addition, we demonstrate that lithium has potential therapeutic targets in casein kinases and calmodulin, suggesting that novel treatment strategies may be possible through these mechanisms.

## Discussion

The present study utilized and validated a midbrain organoid model from CTR and BD patients-derived hiPSCs to explore cellular and molecular phenotypes associated with the disorder. Organoid generation is typically a manual process, labor-intensive, and prone to variability due to researcher handling differences or human error, leading to differences in cell composition and limiting the scalability and reproducibility of results^44,45^. As an alternative, implementing automation could help address some of these challenges, accelerating organoid production and improving the reliability of differentiation outcomes^46^. Therefore, by establishing quality control metrics, implementing liquid handling techniques, and high-content imaging, we aimed to reduce variability and enhance the throughput of organoid cultures, while identifying individual organoids and organoid batches potentially not suitable for downstream experiments. While there were no significant differences in gross morphology or cell type distributions between BD and CTR organoids, electrophysiologically, BD organoids exhibited increased neuronal function, evident through enhanced mean amplitude, conduction velocity and axon length in individual neurons, when compared to CTR. Transcriptome and proteome analyses revealed a compensatory gene regulation network, including pathways associated with calcium signaling and kinase activity. The phosphoproteome analysis indicated significant dysregulation of key signaling pathways in BD organoids, including the PI system, which was partially normalized following lithium treatment.

The observation of heightened neural activity in BD organoids aligns with previous reports from *in vitro* studies of hyperexcitability in psychiatric conditions and elucidates the disorder’s neurobiological underpinnings in a more physiologically relevant 3D environment^7,12^. The midbrain, particularly the ventral tegmental area (VTA), is a crucial dopaminergic site, and alterations in neural activity within this region might contribute significantly to the cyclical switch between manic and depressive states in BD^47–49^. Our data suggest that the heightened functionality of dopaminergic neurons observed in BD organoids could reflect the dysregulation of dopamine signaling and metabolism found in BD patients. This dysregulation in the VTA, along with its connections to other brain regions such as the amygdala, has been implicated in the etiology of the disorder ^49,50^. Thus, our organoid model not only mirrors dopaminergic dysfunctions associated with BD but also offers a potential platform for investigating the underlying mechanisms and testing therapeutic interventions targeting dopaminergic pathways.

A midbrain organoid model represents a significant advancement over traditional monolayer cultures, as it more accurately recapitulates the complex architecture and cellular diversity of the human brain tissue^6,45^. Neuronal subtype analysis of midbrain organoids showed that dopaminergic neurons make up 60-70% of the population, with cholinergic neurons accounting for 35-40%. Glutamatergic and GABAergic neurons each comprising about 1-2%, while astrocytes consistently represented approximately 2% of the cells in both control and BD. This cellular composition aligns with existing literature underscoring the role of excitatory and inhibitory neurotransmitters, such as acetylcholine, glutamate, and GABA, in modulating midbrain dopaminergic neurons^51,52^. The tonic and phasic activity of dopaminergic neurons is critically dependent on cholinergic projections from the hindbrain pedunculopontine and laterodorsal tegmental nuclei. In conjunction with glutamatergic and GABAergic activity, they stimulate nicotinic and muscarinic acetylcholine receptors in the substantia nigra and VTA, modulating dopamine transmission in the dorsal/ventral striatum and prefrontal cortex, effecting processes like arousal, reward, and sleep^52,53^. Our findings of a cholinergic population representing a significant portion of our organoid model highlight the potential for this system to study the modulation of dopaminergic activity. This is particularly relevant given evidence suggesting that increased cholinergic functioning underlies depression, while heightened activity of catecholamines (dopamine and norepinephrine) is associated with mania^54^. Our organoid model, with its balanced representation of dopaminergic and cholinergic neurons, provides a promising platform to further explore and elucidate the complex interplay between these neurotransmitter systems, potentially contributing to more targeted therapeutic strategies for BD.

Previous research in the development of mood disorders has implicated dysregulation of signaling pathways, including those involving brain-derived neurotrophic factor, extracellular signal-regulated kinases, and calcium signaling^13,55,56^. The organoid model enables the study of network-level phenomena and the integration of complex signaling pathways that cannot be replicated in simpler *in vitro* systems. A multi-modal approach integrating electrophysiology, transcriptomics, proteomics, and phosphoproteomics provides a comprehensive understanding of BD pathology, with a focus on midbrain dysregulation. We detected compensatory changes at the transcript and protein level, which not only underscores the importance of intracellular signaling in mood disorders, but may also point toward new therapeutic targets. We hypothesized that BD midbrain organoids may be characterized by specific alterations in the phosphorylation patterns of key proteins involved in these pathways. Notably, proteome and phosphoproteome analyses showed significant dysregulation of BD relevant signaling pathways including phosphatidylinositol (PI), glycogen synthase kinase-3 beta (GSK3B) and protein kinase B (AKT) signaling, validating the relevance of midbrain organoids for studying BD^29,34,35,41,57,58^. We observed dysregulation of CSNK1A1, CSNK2A1, and CALM3 in BD organoids, which was reversed by lithium treatment. This indicates potential molecular mechanisms linking these genes to BD pathophysiology. CSNK1A1 and CSNK2A1 are key regulators of circadian rhythms, inflammation and cellular signaling and play crucial roles in maintaining neurotransmitter balance, which is central to mood regulation and often disrupted in BD^59–63^. CALM3 is a calcium-binding messenger protein that mediates many calcium-dependent processes. Disruptions in calcium signaling are thought to be a feature of BD, affecting neuronal excitability and synaptic plasticity^64^. Calmodulin interacts with various ion channels and neurotransmitter receptors, modulating their activity^64^. Calmodulin also acts as a regulator of calcium/calmodulin-dependent protein kinases (CaMKs) and phosphatases like calcineurin and CaMKKII, which have been implicated in BD and might be directly affected by CALM3^27,65,66^. Future research could focus on further exploring the role of CSNK2A1 and CALM3 in the pathophysiology of BD by examining their interactions with other proteins in the signaling pathways.

Altogether, our work establishes a scalable and functionally validated midbrain organoid platform that offers new opportunities for mechanistic discovery and therapeutic development in bipolar disorder.

## Acknowledgments

We thank members of the larger BD^2^ team for suggestions and discussion, and the Microscopy Resources (MicRoN) core facility at Harvard Medical School for assistance. Some schematics in this paper were created using BioRender.com. This research was funded in whole by BD^2^: Breakthrough Discoveries for thriving with Bipolar Disorder with the grant ID: DG230420.

## Author contributions

K.M., M.W., M.Y, M.C.B.G., P.F., E.Z. performed experiments. H.A.J and M.G.C. assisted in organoid growth, maintenance and sample preparation. M.L. and B.B. performed proteomics and S.M., assisted by N.L. performed bioinformatic analysis of omics data. K.M conceived and co-directed the project, designed overall experimental and analytical strategies, and wrote the manuscript. J.M.T. co-directed the project with K.M., and G.M.C provided significant discussion and consultation of the study.

## Data and materials availability

Supplementary information is made available with this publication. RNAseq data will be deposited at the Gene Expression Omnibus data repository and available upon publication. Raw proteomics data will be deposited in Welcome to MassIVE database and made available upon publication.

## Materials and Methods

### Human induced pluripotent stem cells (hiPSCs)

hiPSCs were generated from human fibroblast cells obtained from different sites. Details of hiPSC lines are listed in Table 1.

**Table 1:** Human induced pluripotent cell (hiPSC) lines used in this study.

| WYSS ID # | Cell Line Origin | Origin ID | Age | Sex | Diagnosis |
| --- | --- | --- | --- | --- | --- |
| **CF1**<sup>1</sup> | Meyer et al<sup>7</sup> | PSC-01-024 | 36 | F | CONTROL |
| **CF2**<sup>2</sup> | Meyer et al<sup>7</sup> | PSC-01-026 | 24 | F | CONTROL |
| **CF3**<sup>2</sup> | NIMH Study ID 163 | MH0185865 | 24 | F | CONTROL |
| **CF4**<sup>1</sup> | NIMH Study ID 163 | MH0185863 | 36 | F | CONTROL |
| **CF5** | Cedars Sinai | CS9BP3iCTR-n1 | 48 | F | CONTROL |
| **CM2** | Meyer et al<sup>7</sup> | PSC-01-020 | 38 | M | CONTROL |
| **CM4** | NIMH Study ID 163 | MH0185932 | 19 | M | CONTROL |
| **CM6** | Cedars Sinai | CS0YX7iCTR-n1 | 49 | M | CONTROL |
| **BF1** | Meyer et al<sup>7</sup> | BD-220-5 | 49 | F | BP-I |
| **BF2** | Meyer et al<sup>7</sup> | BD-81-1 | 35 | F | BP-I |
| **BF3** | Meyer et al<sup>7</sup> | BD-12-33 | 35 | F | BP-I |
| **BF4** | Meyer et al<sup>7</sup> | BD-193-3 | 32 | F | BP-I |
| **BF5** | Meyer et al<sup>7</sup> | BD-PSC-01-002 | 46 | F | BP-I |
| **BF6** | NIMH Study ID 163 | MH0185855 | 22 | F | BP-I |
| **BM2** | Meyer et al<sup>7</sup> | BD-PSC-01-003 | 51 | M | BP-I |
| **BM4** | NIMH Study ID 163 | MH0185869 | 26 | M | BP-I |

<sup>1,2</sup>: May have originated from the same donor, hiPSC lines independently reprogrammed

### hiPSC Culture

hiPSCs were cultured feeder-free on Geltrex LDEV-Free Reduced Growth Factor Basement Membrane Matrix (Gibco) coated plates in mTeSR™1 or mTeSR™ Plus Basal Medium (Stem Cell Technologies). The iPSC lines initially underwent stringent quality control to confirm pluripotency and a normal karyotype. This included alkaline phosphatase assay, karyotype analysis and differentiation into three germ layers.

### Semi-automated Generation of Midbrain Organoids

For embryoid body formation, hiPSCs at 70-80% confluency were used. Cells were incubated with 10 μM ROCK inhibitor (Y-27632, Selleckchem) for 1 hour at 37°C before addition of Accutase to promote detachment. The dissociation process was monitored every 5 minutes, using gentle tapping to aid in breaking up cell clusters, while allowing for some clumping. Once dissociated, cells were quenched by adding 2 mL of mTesr1 medium containing 50 μM ROCK inhibitor before transferring to a 15-mL or 50-mL Eppendorf tube. The cell suspension was centrifuged at 1,000 x *g* for 3-5 minutes to pellet the cells. The supernatant was aspirated, and the cell pellet was resuspended in 1 mL of mTesr1 with 50 μM ROCK inhibitor. Subsequently, cells were counted using a Countess slide (Thermo Fisher Scientific) with Trypan blue exclusion, targeting an ideal cell density of at least 2 × 10^6 cells/mL. Based on cell counts, the suspension was diluted with mTesr1 containing 50 μM ROCK inhibitor to achieve a concentration of 9,000 cells per well of a ULA 96-well plate. The plates were centrifuged at 1,000 x *g* for 1 minute to ensure spherical shaping of embryoid bodies, verified visually, before incubation. For the first two days, the medium in each well was replaced completely, removing as much media as possible without disturbing the embryoid bodies. A new medium, referred to as Midbrain 1st media, was added in a volume of 100 μL per well after aspirating 75 μL. On day 3, a half-medium change was conducted using Midbrain 1st media, omitting the ROCK inhibitor. Day 4 involved a full medium change with additional Midbrain 1st media (E6 medium (Gibco/Fisher Scientific), 1X GlutaMax (Thermo Fisher), 1X MEM-NEAA (Gibco/Fisher Scientific), 1X Penicillin/Streptomycin (Thermo Fisher), 0.055 μM 2-Mercaptoethanol (Sigma Aldrich), 2 μM Puromorphamine (Stemcell Technologies), 10 μM SB-431542 (Stemcell Technologies), 100 ng/mL FGF8 (Stemcell Technologies), and 100 nM LDN-193189 (Stemcell Technologies)). The media was fully replaced with Midbrain 2^nd^ media (DMEM/F-12 (Gibco/Fisher Scientific), 1X N2 Supplement (Thermo Fisher), 1X GlutaMax (Gibco/Fisher Scientific), 1X Penicillin/Streptomycin (Thermo Fisher), 100 nM LDN-193189 (Stemcell Technologies), 100 ng/mL SAG (Cayman Chemical), 2 μM Puromorphamine (Stemcell Technologies), 100 ng/mL FGF8 (Peprotech), 3 μM CHIR-99021 (Stemcell Technologies/Cell Signaling)) at days 6 and 7. Full medium changes were performed on days 8, 10, and 12 using Midbrain 3rd media DMEM/F-12 (Gibco/Fisher Scientific), 1X N2 Supplement (Thermo Fisher), 1X MEM-NEAA (Gibco/Fisher Scientific), 1X GlutaMax (Thermo Fisher), 1X Penicillin/Streptomycin (Gibco/Fisher Scientific), 100 nM LDN-193189 (Stemcell Technologies), 3 μM CHIR-99021 (Stemcell Technologies)). Starting at day 14, media changes took place every other day using Midbrain 4th media Neurobasal medium (Gibco/Fisher Scientific), 1X B27 Supplement (Thermo Fisher), 1X MEM-NEAA (Gibco/Fisher Scientific), 1X GlutaMax (Thermo Fisher), 1X Penicillin/Streptomycin (Gibco/Fisher Scientific), 200 μM Ascorbic Acid (Sigma Aldrich), 500 μM cAMP (Sigma Aldrich), 20 ng/mL BDNF (Peprotech/Thermo Fisher), 20 ng/mL GDNF (Peprotech/Thermo Fisher), and 1 ng/mL TGF beta (Stemcell Technologies).

#### Automated media changes using Fluent 780 (Tecan)

To enable high-throughput maintenance of brain organoids, we optimized media changes for embryoid bodies and organoids using Tecan’s Fluent 780 liquid handling technology. We developed a dynamic media change protocol that allows for real-time adjustments in media removal and addition, as well as the number of plates needing media change. The system incorporates liquid level sensing with Tecan’s Flexible Channel Arm (FCA). Before media removal, the FCA attaches a 1000 µL tip (Tecan, Part #: 30000631) which measures the meniscus height in the A1 well of each plate. This data is then communicated to the Multi-Channel Arm (MCA), equipped with a 96-air displacement unit and 150 µL filtered tips (Tecan, Part #: 30038618), for media aspiration. The protocol calculates meniscus height using user-defined plate specifications and incorporates a 0.5 mm buffer to prevent organoid aspiration. Aspiration speed is maintained at 1.88 µL/second. Once media removal is completed across all plates, fresh media is added using a new set of tips at a rate of 20 µL/second.

#### Imaging and Analysis of Organoid Development Using Ramona’s Vireo System

For high-throughput imaging, we employed the Ramona Vireo™ system and MCAM^TM^ software (Ramona Optics), which is equipped with 24 cameras enabling both widefield and fluorescence imaging at a fast rate. Using the MCAM™ software, we established a detailed protocol specifying parameters such as cell batch placement, optic type, exposure, intensity, and imaging z-height. Specifically, the optic type was set to High-Contrast Brightfield, with an exposure setting of 35, an intensity of 40, and a z-height of 10.02 mm. Within the MCAM^TM^ software, a plate map was defined, and a corresponding spreadsheet was generated to document the defined names and their associated well locations. Images were automatically stored in folders labeled with the date in a DDMMMYYYY format (e.g., 12FEB2025) to ensure systematic data management. Each plate was rapidly imaged, taking only a few seconds per plate. The resulting data was then segmented within the MCAM^TM^ software, providing measurements of organoids area, eccentricity and circularity. The segmented data was subsequently processed using MATLAB, which compiled all segmentation analyses with metadata including Cell-ID, Days-In-Culture, Plate Number, Batch-ID, and Disease State. For further analysis and visualization, the processed data were imported into GraphPad Prism.

#### Treatment of Midbrain Organoids

Lithium carbonate treatment was performed at a concentration of 100 μM for a duration of one week, starting at DIV46, with the medium containing the drug being replenished every other day. This concentration reflects levels typically observed in the cerebrospinal fluid of patients with bipolar disorder who are undergoing lithium treatment^67^.

### Microelectrode Array (MEA) using MaxTwo (MaxWell Biosystems)

#### Surface Sterilization and Pre-conditioning of electrode array

To ensure optimal organoid adherence and function, MaxTwo 6-Well Plates underwent surface sterilization and pre-conditioning. Initially, surface preparation was achieved using 2 mL per well of a 1% Terg-a-zyme solution followed by 2 hours of incubation at room temperature. After incubation, the Terg-a-zyme solution was removed, and each well was thoroughly washed three times with deionized water to ensure complete removal of the cleaning solution. Following surface preparation, plate sterilization was carried out by thoroughly spraying the plates with 70% ethanol on both front and back surfaces. Each well and compartment were filled with ethanol to ensure effective sterilization. After 30 minutes in a biological safety cabinet, the ethanol was aspirated, and wells were washed three times with sterile deionized water. The plates were then completely dried using a vacuum pump. For medium pre-conditioning, each well was filled with 1.2 mL of culture medium. The plates were placed in a 37°C, 5% CO2 incubator for 2 days.

#### Surface Coating

For primary coating a 0.07% PEI (Polyethyleneimine) solution was prepared by diluting a 7% PEI stock solution with a 1X borate buffer. This PEI solution was filtered through a 0.22 μm filter for sterilization. Each well received 50 μL of the solution, covering the electrode arrays, followed by incubation at 37°C, 5% CO2 for 1 hour. After incubation, the coating solution was aspirated, and wells were washed. For the secondary coating, laminin was thawed on ice and diluted to the working concentration in chilled culture medium. Each well was treated with 50 μL of the secondary coating solution and incubated overnight under the same incubation conditions to promote effective laminin binding.

#### Organoid Plating

Before transferring organoids, the wells were prepared by carefully aspirating the secondary coating solution, leaving a thin film. On DIV36 three, whole organoids were positioned at the center of the electrode array using small droplets of fresh Midbrain 4th media, applied via pipetting or with the assistance of a fine sterile brush to ensure precise placement. Excess medium was removed, allowing the surface to dry and enhance adherence. Organoids were given 1–2 minutes to adhere before gently adding 50 μL of culture medium. This procedure was iterated across the necessary wells, and plates were incubated for 2 hours in a CO2 incubator. Subsequently, each well was filled with fresh medium, ensuring the organoids remained hydrated. By day 41, a half-medium change was performed on MEA wells using BrainPhys media (StemCell Technologies), carefully avoiding tilting the plate or touching the electrodes with the pipette tip during aspiration. The media added the previous day was allowed to equilibrate, allowing for the first baseline activity scan to be conducted on day 42 using the MaxTwo MEA system. Subsequent half-medium changes in MEA wells with BrainPhys media were performed on days 43 and 45, followed by the second and third baseline activity scans on days 44 and 46, respectively. From day 48 to day 88, half-medium changes with BrainPhys media were carried out every other day. MEA recordings were performed between media changes to monitor and record neural activity or axon tracking.

#### Neural Activity Recording

Organoid cultures were maintained at 37°C in a 5% CO2 incubator. The data acquisition for the Activity Scan Assay was configured with “Sparse 7x,” and recordings were set for 30 seconds per configuration. The “Record Only Spikes” option was active, and real-time monitoring was facilitated by enabling views for both “Electrodes” and “Traces.” For the Axon Tracking Assay, recording time was set to 180 seconds. Pre-assay settings included a 2.5 ms window before and a 5 ms window after axon potential detection. Density was set for full electrode coverage, with either “Array Section” or “Blocks” mode based on coverage needs. Custom boundaries could be defined using screen interface controls. Analysis parameters for the activity assay included a firing rate threshold of 0.10 Hz, amplitude threshold of 20.00 μV, and an ISI threshold of 200 ms. For axon tracking, analysis parameters included a spike threshold of 20, footprint completeness threshold of 0.75, enabled plot branch level, a latency threshold set to 0.00 ms, and a radius of 3 pixels. Following analysis, data were exported to external devices using the system’s export functionality, ensuring all metrics were reliably secured for consistent assessment and replication. This comprehensive setup and protocol guaranteed high-quality data acquisition and analysis of neural activity in the plated organoids.

### Transcriptomics

#### RNA Extraction from Midbrain Organoids

A total of 5-8 organoids were first lysed in lysis buffer from the RNeasy Mini Kit (Qiagen) by pipetting. The lysate was then passed through QIAshredder (Qiagen) columns prior to following the RNeasy Mini Kit manufacturer’s recommended protocol including a DNAse digestion step.

#### RNA sequencing read alignment and quantification of gene expression

RNA quality control, library preparation and sequencing were facilitated by Novogene Corporation. RNA sequencing was performed using the NovaSeq 6000 System with sequencing of 20 million paired-end 150 bp reads. For organoids, reads were aligned to GRCh38 with Ensembl GRCh38.92 gene models using STAR version 2.5.4a with options -- alignSJDBoverhangMin 1 --alignSJoverhangMin 8 --outFilterMultimapNmax 20 –outFilterType BySJout --alignIntronMin 20 --alignIntronMax 5000000 --alignMatesGapMax 5000000. Expression levels of genes were quantified as gene counts using STAR version 2.5.4a with option --twopassMode Basic.

#### Gene annotation

Gene annotation and gene sets used for functional gene classification into biological process (BP), molecular function (MF), and cellular component (CC) were from the GRCh38.p12 database downloaded from Ensembl Biomart on March 4, 2018.

### Proteomics

#### Individual Brain Cell Specific Sample Preparation

Samples were undergoing an S-trap protocol (Protifi, NY) procedure to break cells and open proteins from their 3D structures for digestion (1). Digestion: A stock solution of Trypsin Platinum, Mass Spec Grade (Promega) was made up at 100 µg/mL in 50 mM TEAB. A specific volume of stock trypsin was added to each sample vial for each set of brain cells using a 1:50 ratio (Trypsin:Protein). Samples were incubated for 2 hours at 50 °C in S-trap midi columns on Eppendorf ThermoMixer. Digested samples were then transferred to HPLC glass vials and were resuspended in 6 µL of 0.1% Formic Acid in ultrapure HPLC grade water and were then injected on LC-MS/MS.

#### Phosphoproteomics enrichment procedure

The phosphoproteomics was performed at Harvard Center for Mass Spectrometry (HCMS). S-trap mini (Protifi, NY) according to manufacturer protocol. Briefly, samples were resolubilized in 5% SDS for reduction and alkylation, further digested in buffer trypsin (Promega) for 5 hours. The digested samples were enriched by High-Select™ TiO2 Phosphopeptide Enrichment Kit (Thermo-Fisher) according to the vendor’s instructions. Enriched phosphopeptides were labeled with TMT18plexPRO (Thermo-Fisher) according to manufacturer protocol. Sample fractions were submitted for single LC-MS/MS experiment that was performed on Exploris 240 Orbitrap (Thermo, MA)

#### Mass spectrometry analysis

After separation each fraction was submitted for single LC-MS/MS experiment that was performed on a Exploris 240 Orbitrap (Thermo Scientific) equipped with NEO ( Thermo Scientific) nanoHPLC pump. Peptides were separated onto a 75 µmx4 cm C18 trapping column (Premier LC, CA) followed by analytical column PepMap Neo 50umx150mm (Thermo Scientific, Lithuania). Separation was achieved through applying a gradient from 5–25% ACN in 0.1% formic acid over 60 min at 200 nl min^−1^. Electrospray ionization was enabled through applying a voltage of 2.1 kV using a PepSep electrode junction at the end of the analytical column and sprayed from stainless still PepSep emitter SS 30 µm LJ (Bruker, MA). The Exploris Orbitrap was operated in data-dependent mode for the mass spectrometry methods. The mass spectrometry survey scan was performed in the Exploris 240 Orbitrap in the range of 450 –900 m/z at a resolution of 1.2 × 10^5^, followed by the selection of the ten most intense ions (TOP10) ions were subjected to HCD MS2 event. The fragment ion isolation width was set to 0.8 m/z, AGC was set to 50,000, the maximum ion time was 150 ms, normalized collision energy was set to 34V and an activation time of 1 ms for each HCD MS2 scan.

#### Mass spectrometry data analysis

Raw data were submitted for analysis in Proteome Discoverer 3.1.1.93 (Thermo Scientific) software with Chimerys. Assignment of MS/MS spectra was performed using the Sequest HT algorithm and Chimerys (MSAID, Germany) by searching the data against a protein sequence database including all entries from the Mouse Uniprot database (SwissProt 19,768 2019) and other known contaminants such as human keratins and common lab contaminants. Sequest HT searches were performed using a 20 ppm precursor ion tolerance and requiring each peptides N-/C termini to adhere with Trypsin protease specificity, while allowing up to two missed cleavages. Carbamidomethyl on cysteine amino acids (+57.021464 Da) and TMT18plexPRO modification on Lysine’s and peptide N-terminus were set as permanent modifications while methionine oxidation (+15.99492 Da) was set as variable modification. A MS2 spectra assignment false discovery rate (FDR) of 1% on protein level was achieved by applying the target-decoy database search. Filtering was performed using a Percolator (64bit version)^68^.

### Cell Type Quantification using FACS

#### Organoid Dissociation and Cell Fixation

Organoids were initially washed twice in each well using phosphate-buffered saline (PBS) to ensure the removal of culture media. For enzymatic dissociation, the organoids were incubated with 300-500 μL of Accutase for 15 minutes at room temperature. Subsequent mechanical dissociation was performed by repeated pipetting to achieve a single-cell suspension.

After dissociation, cells were centrifuged at 1,000 x *g* for 3 minutes to pellet the cells. The supernatant was discarded, and the cell pellet was subjected to a wash with cold PBS. To initiate fixation, the cells were dissociated and resuspended in 500 μL of cold 4% paraformaldehyde (PFA) and incubated on ice for 1 hour to ensure adequate fixation. Following fixation, cells were centrifuged at 3,000 x *g* for 1 minute. The supernatant was carefully removed, and the cells were washed three times with 1 mL PBS, with each resuspension involving vortexing to ensure thorough washing. Fixed cells were stored in PBS until the permeabilization process.

#### Immunocytochemistry (ICC) Staining Procedure

To begin the ICC staining process, cells were recovered by centrifugation at 3,000 x *g* for 1 minute, and cell counting was performed to ensure a minimum of 100,000 cells per sample. Following this, the supernatant was removed, and the cell pellet was resuspended in 300 μL of blocking buffer composed of 3% Normal donkey serum and 0.3% Triton X-100 in PBS. The cells were incubated in this blocking buffer for 30 minutes at room temperature, followed by brief vortexing at 30% speed. Subsequently, the cells were centrifuged at 3,000 x *g* for 1 minute, and the supernatant was discarded. The cell pellet was resuspended in 100-250 μL of blocking buffer containing the primary antibodies, with vortexing at 70% speed to ensure uniform antibody distribution. The tubes were placed in a rack holder and secured with tape. The samples were incubated overnight on a shaker at 4°C to allow for proper antibody binding.

After the primary antibody incubation, the cells were washed three times with 500 μL PBS, with each wash followed by centrifugation at 3,000 x *g* for 1 minute to remove unbound antibodies. The cells were then resuspended in 100-250 μL of blocking buffer containing the secondary antibodies and incubated at room temperature for 1 hour, with brief vortexing at 30% speed.

Finally, the cells were washed three times with 500 μL PBS, centrifuging at 3,000 x *g* for 1 minute each time, to remove excess secondary antibodies. The cells were resuspended in 0.5-1 mL PBS and passed through a 0.45 μm mesh to eliminate any aggregates. The stained cells were kept on ice and protected from light until flow cytometric analysis.

| Target | Host | ID | Company |
| --- | --- | --- | --- |
| βIII-tubulin (Tuj1) | Mouse | 5568S | Cell Signaling Technologies |
| Tyrosine Hydroxylase (TH) | Sheep | NB300-11055 | Novus Biologicals |
| GABA | Rabbit | A2052 | Millipore-Sigma |
| GFAP | Mouse | 3670S | Cell Signaling Technologies |
| VGlut1 | Mouse | AB242204 | Abcam |
| Choline Acetyltransferase (ChAT) | Goat | AB144P | Millipore-Sigma |

#### FACS and Cell Type Quantification

Cell sorting was conducted using the Sony SH800 flow cytometer (Sony Biotechnology). Initial settings were established by adjusting the voltages using unstained control samples to optimize signal detection. The data were presented on a logarithmic scale, and the threshold was decreased to enhance visualization of the target cell population in the forward scatter (FSC) versus side scatter (SSC) plot. Selection of single cells and subsequent gating was informed by negative controls, employing the following gating strategies to identify subpopulations: ßIII-Tubulin^+^/TH^+^, ßIII-Tubulin^+^/ChAT^+^, ßIII-Tubulin^+^/GABA^+^, GFAP^+^, and VGlut1^+^. Prior to sorting, the Sony SH800 was precisely calibrated according to the manufacturer’s guidelines. Lasers and optical filters used on the Sony SH800 included: Violet laser (405 nm) with a 450/50 nm bandpass filter for collecting BV421 signals. Blue laser (488 nm) with a 525/50 nm bandpass filter for Alexa Fluor 488 (AF488) or GFP signals. Yellow laser (561 nm) with a 600/60 nm bandpass filter for Alexa Fluor 594 (AF594) or PE signals. Red laser (647 nm) with a 665/30 nm bandpass filter for Alexa Fluor 647 (AF647) or APC signals. Subsequent analysis was performed on a FACSymphony A3 (BD Biosciences). Quality control of the FACSymphony A3 was conducted following the manufacturer’s recommendations. Voltages and gating parameters on the A3 were established using control samples to ensure accurate signal discrimination. The lasers and optical filters utilized on the FACSymphony A3 comprised: a. blue laser (488 nm) with a 512/20 nm bandpass filter for detecting Alexa Fluor 488 emission; b. yellow laser (561 nm) with a 585/15 nm bandpass filter for capturing Alexa Fluor 594 emission; c. red laser (640 nm) with a 670/30 nm bandpass filter for acquiring Alexa Fluor 647 emission. This approach ensured precise gating, accurate cell population identification, and reliable analysis of the target fluorescence markers.

### Immunofluorescence Microscopy

Organoids were fixed by first removing their culture media and then incubating them with 4% paraformaldehyde (PFA) at 37°C for 5 minutes. The organoids were subsequently washed once with phosphate-buffered saline (PBS), being cautious to avoid further washing to prevent detachment of neuron cells. Following fixation, organoids were stored at 4°C in PBS until they were ready to be stained. To reduce background fluorescence and inhibit residual fixation activity, organoids were quenched using a 0.1M glycine solution in PBS for 15 minutes at room temperature. Organoids were incubated in a permeabilization buffer consisting of 3% normal goat serum (NGS) or normal donkey serum (NDS) and 0.3% Triton X-100, dissolved in 10 mM PBS, for a duration of 20-60 minutes. The choice of serum was contingent on the specificity of the secondary antibody used. The primary antibody was prepared by centrifugation in a solution of 3% NGS or NDS and 0.2% Triton X-100 at maximum speed for 5 minutes. The organoids were then incubated with the primary antibody solution for 1-2 hours at room temperature or alternatively overnight at 4°C in a humidified chamber. Post-incubation, the organoids were washed three times for 10 minutes each with PBS. Secondary antibody preparation involved centrifugation under the same conditions as the primary antibody in 3% NGS or NDS and 0.2% Triton X-100 for 5 minutes at maximum speed. Organoids were incubated with the secondary antibody in a humidified and dark chamber for 30 minutes to 2 hours. Following exposure, specimens underwent three successive 10-minute PBS washes. For sample preservation and imaging, organoids were mounted by applying one drop of mounting medium, such as ProLong Gold, onto a microscope slide and covered with a cover slip. Samples were left to dry overnight in a dark, dry environment, such as a cardboard box, which effectively absorbs moisture and protects fluorescence.

## Figure Legends

**Figure S1:**
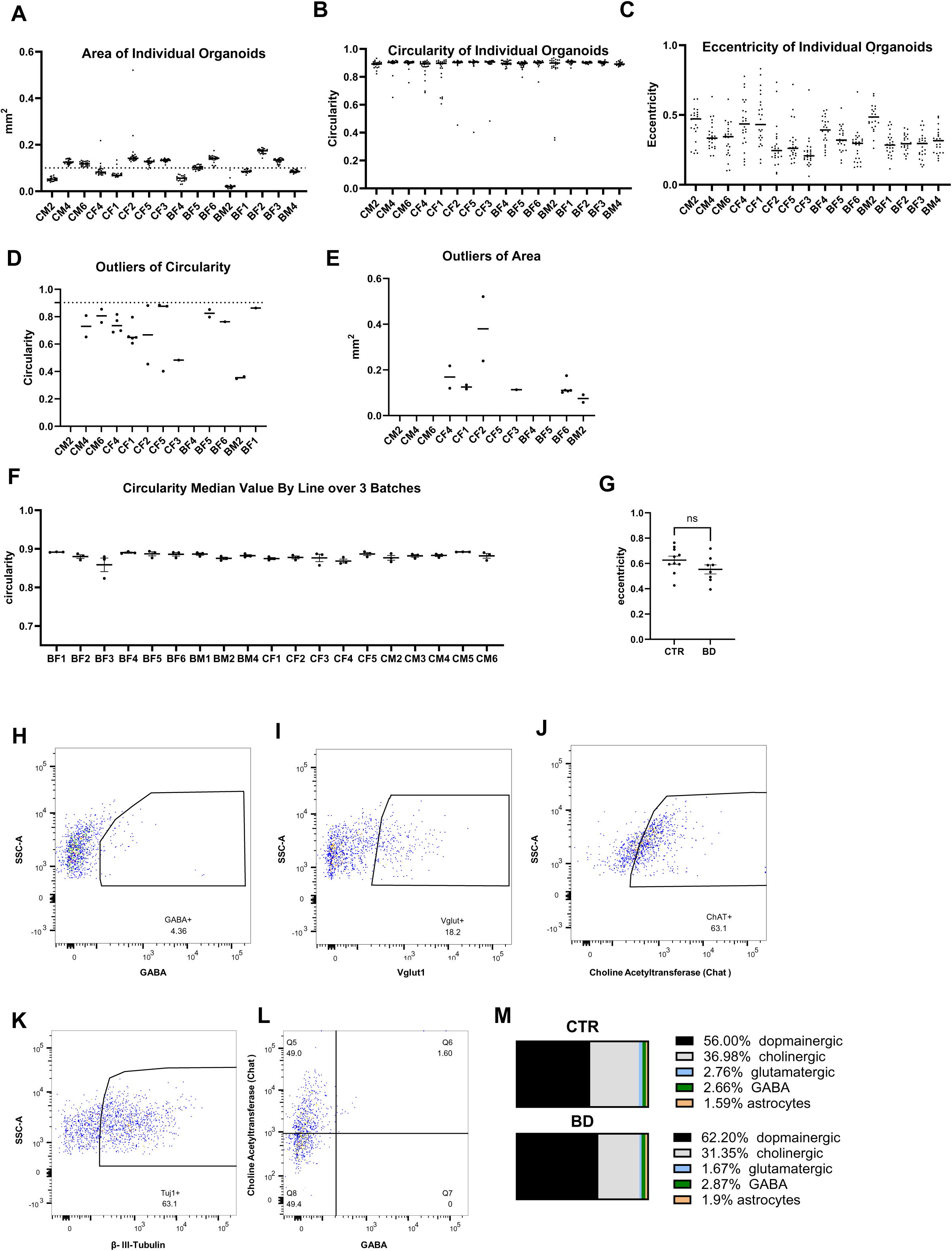
Additional organoid measurements and FACS gating strategies: **A-C** Area (A), circularity (B) and eccentricity (C) of 24 individual organoids from control (CTR, n=8) and bipolar disorder (BD, n=8) lines. **D:** Outliers of circularity identified by ROUT outlier analysis (Q=5%) of individual organoid level data in (B). **E:** Outliers of area identified by ROUT outlier analysis (Q=5%) of individual organoid level data in (A). **F:** Mean circularity of CTR n=8 and BD n=8 lines over 3 technical organoid batches. **G:** Mean eccentricity in DIV35 midbrain organoids. Each datapoint represents the mean of 24 individual organoids from each iPSC line. **H-L**: FACS sorting gating strategies for GABA, Vglut and ChAt. **M:** Cell type distribution of in CTR and BD organoids.

